# Chromosomes remain individualized through interphase in embryos of the tardigrade *Hypsibius exemplaris*

**DOI:** 10.1101/2025.07.25.666853

**Authors:** Lillian D. Papell, Adriana N. Coke, Bailey N. de Jesus, Clayton J. Harry, Pu Zhang, Bob Goldstein

## Abstract

Tardigrades are microscopic animals that can survive exceptional levels of ionizing radiation or desiccation – DNA-damaging conditions that would kill most animals. Irradiation or radiomimetic drug treatment of the tardigrade *Hypsibius exemplaris* can induce remarkably high expression levels of DNA repair genes, primarily those in the base excision repair and nonhomologous end joining pathways. How tardigrades can repair widespread DNA damage without producing frequent, large-scale chromosome structural abnormalities, like chromosome translocations and fusions, is unknown. Here, we report the results of examining chromosome and nuclear architecture throughout the cell cycle in early embryos of *H. exemplaris*. We found that *H. exemplaris* chromosomes are maintained in an individualized form throughout the cell cycle. We were surprised to also find that each chromosome is housed in a fully or partially separate lamin-lined compartment, instead of all chromosomes being housed in a single, nearly spherical nuclear lamina and envelope. Our results reveal unusual chromosomal and nuclear organization in a tardigrade. We speculate that these unexpected features might limit chromosomal rearrangements during DNA damage repair in extreme conditions.

**SIGNIFICANCE STATEMENT:**

- We have investigated unusual chromosome organization in an organism that survives extremely DNA-damaging environments, a tardigrade, through early embryonic cell cycles.
- Chromosomes in fixed and stained embryos appeared more condensed through interphase than is typical for animal cells.
- Chromosomes remained individualized, in fully or partially separate lamin-lined compartments, through interphase.
- The results reveal a unique nuclear and chromosomal organization in tardigrades, which we speculate might contribute to limiting chromosomal structure abnormalities under DNA damaging conditions in nature.

## INTRODUCTION

Tardigrades, or water bears, are renowned for their ability to survive extreme conditions such as ionizing radiation, desiccation, and extreme temperatures (Kinchin, 1994). Some progress has been made in uncovering the mechanisms behind tardigrade resistance to stresses (see Hashimoto and Kunieda, 2017; Hibshman et al., 2020; Arakawa, 2022 for review). For example, irradiation causes massive DNA damage in *H. exemplaris*, but the animals repair the damage and survive (Clark-Hachtel *et al*., 2024; Anoud *et al*., 2024). For perspective, the median lethal dose (LD50) for ionizing radiation in humans is 5 Gy, while the LD50 for *H. exemplaris* is between 100 Gy and 500 Gy for embryos, and ∼4,000 Gy for adults (Beltrán-Pardo *et al*., 2015). In response to extremely high levels of ionizing radiation, genes involved in DNA repair mechanisms such as nonhomologous end joining (NHEJ) become dramatically upregulated, with some genes upregulated almost 300-fold in *H. exemplaris* and a closely related species (Clark-Hachtel *et al*., 2024; Anoud *et al*., 2024; Li *et al*., 2024).

Desiccation and irradiation are expected to induce many, simultaneous DNA double strand breaks in tardigrades and other organisms (Dose *et al*., 1992; Steel, 1996; Neumann *et al*., 2009; Clark-Hachtel *et al*., 2024). Many tardigrade species survive desiccation and indeed are likely to experience desiccation in nature (Kinchin, 1994). Repair of multiple double-strand breaks at once, especially through the NHEJ pathway, can result in errors and even large chromosomal rearrangements (Lieber *et al*., 2010). If chromosomal rearrangements occur in germ cell precursors, adult tardigrades would be passing these changes directly to their offspring. Repeated genome reorganization after DNA damage could have drastic evolutionary consequences, but chromosome fusions and translocations appear very rare in a comparison of chromosome-level assemblies from two different *Hypsibius* species (Li *et al*., 2024). It is unknown how *H. exemplaris* endures or fixes DNA damage from these extreme environments without extensively impacting the genome. An understanding of whether tardigrades tolerate these potentially harmful changes or repair damage in a way that avoids chromosomal rearrangements would be useful to inform further studies of tardigrade resistance to extremes.

In an early stage of the work reported here, we made the fortuitous qualitative observation that chromosomes appeared more condensed than expected in some cells of *H. exemplaris* embryos. Embryonic cells of animals typically cycle between an interphase consisting solely of the DNA replication period (S phase), and mitosis (Farrell and O’Farrell, 2014; Kipreos and van den Heuvel, 2019). Chromosomes are fully decondensed during interphase, causing the DNA to appear by light microscopy methods largely as a diffuse sphere filling most of each cell’s nucleus. The chromosomes begin to condense during prophase of mitosis, just before nuclear envelope breakdown, and are most condensed at metaphase and anaphase. In mitosis, the chromosomes also individualize, i.e. take shapes that are separate from one another. These two processes, condensation and individualization, occur by partially separate mechanisms, and specific mechanisms also maintain chromosome individuality once it is initially established (Cuylen *et al*., 2016; Schneider *et al*., 2022; Hibino *et al*., 2024; Raaijmakers *et al*., 2025). At late anaphase and telophase, chromatin decondenses, re-establishing the interphase chromatin state (Antonin and Neumann, 2016). Chromosome condensation and individualization are essential during cell division cycles to ensure that chromosomes are accurately partitioned into the two daughter cells (Koshland and Strunnikov, 1996). The opposite process, chromosome decondensation, is important to allow transcriptional machinery and other proteins to access DNA and control gene expression during interphase (Shoaib et al., 2020).

We set out to investigate nuclear organization throughout the cell cycle in *H. exemplaris* embryos. First, we looked for irregularity in the chromosome condensation cycles by creating a timeline of fixed, DNA-stained embryos throughout the early cell cycles. We also performed imaging with various *in vivo* stains to further understand chromosome condensation state and individualization. Lastly, we looked at overall nuclear lamin organization by performing lamin immunostaining. The findings suggest that chromosomes in *H. exemplaris* embryos are highly atypical in that they remain individualized throughout the cell cycle, isolated into separate or partially-separate nuclear lamin-lined compartments.

## RESULTS

### Chromosomes remain more condensed in *H. exemplaris* early embryonic cell cycles than in a more typical system, *C. elegans*

After noticing unexpectedly condensed chromosomes in some fixed *H. exemplaris* embryos that included interphase cells, we attempted to develop a protocol for live imaging of *H. exemplaris* DNA by testing a large number of DNA dyes (see Methods). Propidium iodide has worked before upon electroporation into embryos (McGreevy *et al*., 2018), and we found by soaking embryos in dyes that NucBlue (Hoechst 33342) worked intermittently, and Kakshine Yellow (formerly called PC1; Uno *et al*., 2021) worked reliably (discussed further below), but we failed to find a dye that would both mark chromatin and allow normal development through live imaging experiments. We also found that both NucBlue and Kakshine Yellow resulted in bright fluorescence of cytoplasmic foci, calling into question whether these dyes labeled only DNA in living *H. exemplaris* embryos, or additionally either RNA or other cellular components.

Therefore, we tested the hypothesis that *H. exemplaris* chromosomes never fully decondense during early embryonic divisions by creating a timeline of fixed, DAPI-stained embryos. Cells of early-stage embryos are large, presenting an opportunity to resolve cytological features well by light microscopy methods. We fixed embryos at 15 minute intervals from 2 to 6 hours after embryos were laid by their mothers (hours post laying, hpl). After the first cell cycle, which takes about 2 hours, each cell cycle is approximately 50 minutes long (diagrammed in Figure 1A). Therefore, the timeline covered the 2 to 32 cell stages (Gabriel *et al*., 2007), with approximately 3-4 timepoints in each cell cycle after the first cell cycle. Embryonic cells in *H. exemplaris* divide in an asynchronous and predictable pattern starting at the 2-cell stage (Gabriel *et al*., 2007), so most nuclei should be in interphase in most of the timepoints we examined, and nearly every timepoint we examined should include some interphase nuclei.

**Figure 1.**
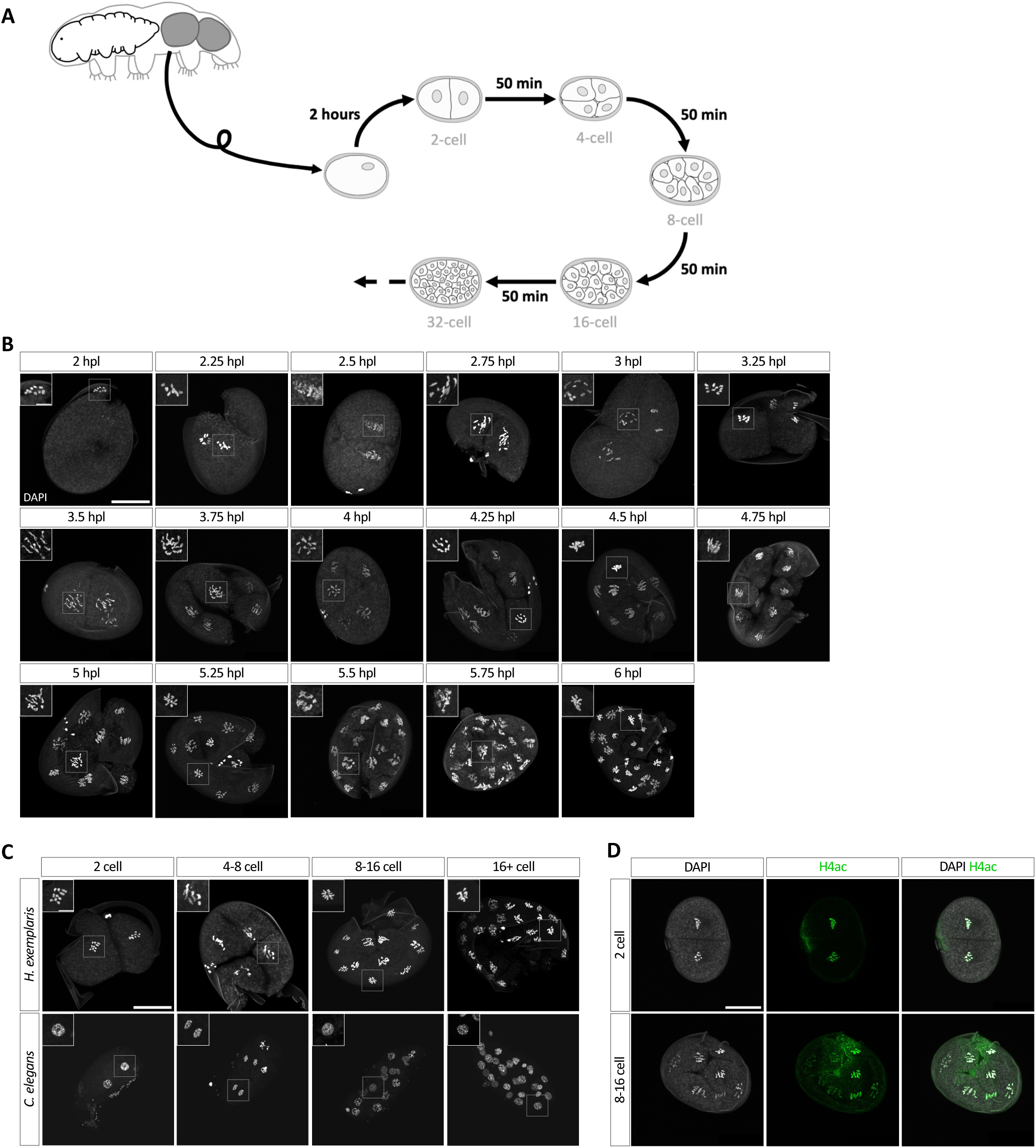
Chromosomes do not fully decondense in *H. exemplaris* early embryonic cell cycles. (A) Schematic diagram of a mother laying embryos in her cuticle and early embryonic development in *H. exemplaris*, including the length of each cell division and corresponding cell stage. (B) Enrichment patterns of DAPI (white) in 2 to 32 cell tardigrade embryos, with zoomed-in views of cells with contrast adjusted in top left corners, including polar body chromosomes at 2 hpl. The timeline spans 4 hours from 2 to 6 hpl, with representative embryos approximately 15 minutes apart. 61 embryos were imaged in total, with an average of 3.6 embryos per time point. (C) Enrichment patterns of DAPI (white) in *C. elegans* and *H. exemplaris* embryos, with zoomed-in views of cells with contrast adjusted in top left corners. Representative embryos range from the 2 to past the 16 cell stage. (D) Localization patterns of DAPI (white) and histone H4ac (green) in representative 2-6 hours post laying (hpl) *H. exemplaris* embryos. Scale bars: 20 µm for whole embryo views, 5 µm for insets.

We observed that in all identifiable nuclei throughout these early embryonic cell cycles, no chromosomes appeared as decondensed as we had originally expected (Figure 1B). To compare what we observed in *H. exemplaris* embryos with more typical cells, we carried out the same fixation and staining protocol in parallel on *C. elegans* embryos. Nematodes (which include *C. elegans*) and tardigrades are both members of the superphylum Ecdysozoa (Aguinaldo *et al*., 1997; Giribet and Edgecombe, 2017). *C. elegans* cells go through condensation and decondensation cycles typical of animal cells and have been used as models for studying cell cycle dynamics and condensation/decondensation mechanisms (see Kipreos and van den Heuvel, 2019 for review). We observed that *C. elegans* chromosomes were less condensed and individualized, as expected for typical interphase nuclei, than the *H. exemplaris* chromosomes: in the *C. elegans* embryos, the DAPI-stained DNA in many of the nuclei appeared as diffuse spheres characteristic of decondensed chromatin in interphase. In contrast, all the *H. exemplaris* nuclei contained distinct, condensed chromosomes (Figure 1C). We quantified this apparent difference in condensation using an assay developed previously (Maddox *et al*., 2006). This assay relies on intensity measurements of fluorescently marked chromatin to assess the degree of chromatin condensation. We normalized fluorescence intensities as in Maddox *et al*., 2006 and measured the percentage of pixels at gray values in the bottom 20% of the intensity range (Figure S1A-B) – a measurement that grows as chromosome condensation progressively clears bright fluorescence away from ever-larger regions of the nucleus (Maddox *et al*., 2006). The results quantitatively confirmed significantly greater condensation on average in *H. exemplaris* nuclei than in *C. elegans* nuclei (Figure S1C).

We next wondered whether this unexpectedly condensed state of chromatin could be detected throughout the *H. exemplaris* life cycle (diagrammed in Figure S1E). We repeated the same fixation and staining protocol in later embryonic stages from the 60 cell stage to hatchling, and in specific adult cells – epidermal, pharyngeal, and intestinal cells – in both *H. exemplaris* and *C. elegans*. Small nuclei found in later embryonic stages and in some adult cells make condensation state difficult to assess, since even condensed chromosomes may fill most of a small nucleus by light microscopy methods. We did not detect a significant difference in chromosome condensation between the two species at later embryonic stages or in pharyngeal or intestinal nuclei, but we could detect significantly greater condensation in *H. exemplaris* epidermal nuclei than in *C. elegans* epidermal nuclei (Figure S1F-I). We conclude that an unexpectedly condensed state of chromosomes exists in early embryos and at least some adult cells.

The DNA stain we used, DAPI, is a fluorescent minor-groove binder that preferentially marks AT-rich DNA (Kapuścínski and Szer, 1979). Because this preference might introduce a bias in what parts of chromosomes fluoresce most brightly, we also stained embryos with an antibody recognizing acetylated histone H4 (H4ac). Acetylation of multiple residues of the histone H4 tail is typical of transcriptionally active chromatin, which is generally decondensed in other organisms (see Dhar *et al*. 2017 for review), and early embryos of *H. exemplaris* have RNA abundance patterns that change rapidly, suggesting active transcription at these stages (Levin et al., 2016). We saw that H4ac immunostaining overlapped with DAPI staining (Figure 1D), and there was no quantitative difference in the degree of condensation between H4ac and DAPI staining (Figure S1D). We conclude that fixed embryonic cells of the tardigrade *H. exemplaris* have chromosomes that are more condensed than in a representative of more typical cells, *C. elegans*.

### Chromosomes remain unusually individualized in *H. exemplaris* early embryonic cell cycles

Our observations of chromosomes in the experiments above also suggested to us that chromosomes remain unusually individualized, i.e. in a state in which most individual chromosomes could be recognized and counted, even in interphase by light microscopy, rather than as a single diffuse mass that filled most of a spherical or nearly spherical nucleus (Figure 1B,C,D). To assess quantitatively how separate the chromosomes were from one another, we used Imaris Machine Learning to segment and count chromosomes (pipeline diagrammed in Figure 2A). *H. exemplaris* is diploid and has 5 pairs of chromosomes (2n = 10 chromosomes) (Gabriel *et al*., 2007). Much as we expected based on direct observations, some chromosomes were too close to each other to be confident that they were individualized, and the software recognized 5-10 chromosome masses per nucleus (Figure 2B). There were few nuclei with greater than 10 chromosomes, which suggests that over-segmentation of chromosomes was limited. These results suggest that at least partial chromosome individualization is maintained in most nuclei. All 10 chromosomes could be counted especially frequently in cells of 4-cell stage embryos, where nuclei are large, compared to later stages with smaller nuclei (Figure S2). We conclude that chromosomes remain unusually individualized in *H. exemplaris* early embryonic cell cycles.

**Figure 2.**
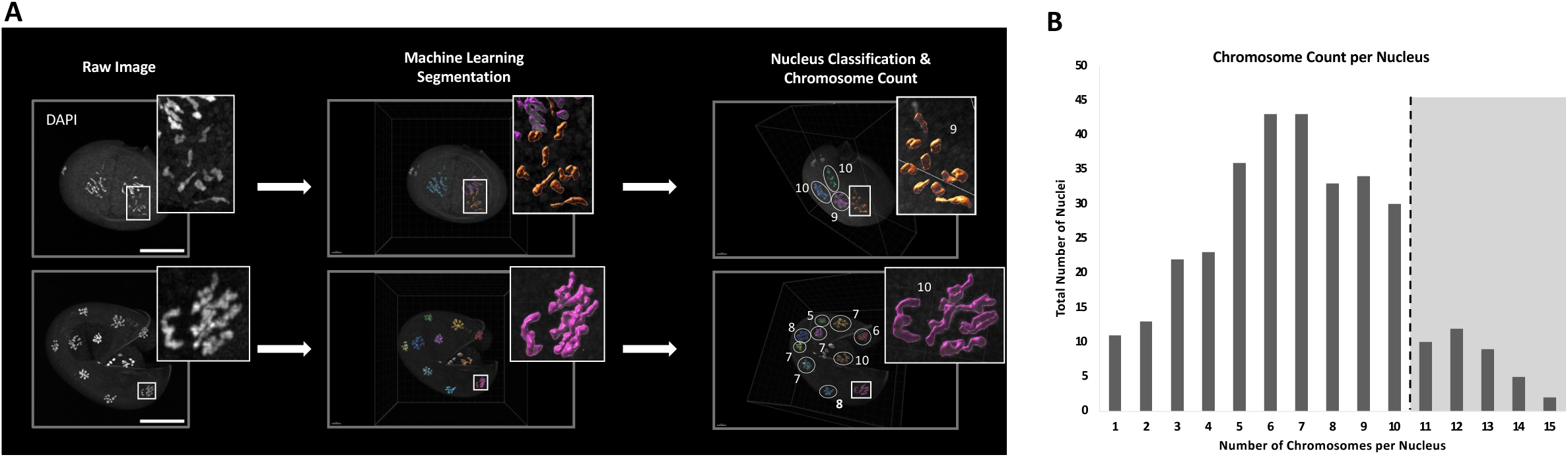
Chromosomes remain individualized in *H. exemplaris* early embryonic cell cycles. (A) Imaris Machine Learning pipeline with representative images. Scale bars: 20 µm. (B) Histogram of the chromosome count per nucleus from Imaris segmentation. The section highlighted is the chromosome count greater than the known number in *H. exemplaris*, 2n = 10 chromosomes.

Since individualized chromosomes are not normally seen in animal cells in interphase, we examined whether individualized chromosomes would be seen using other dyes and in the absence of fixation. Among the dyes we tested above, we used the two dyes that most specifically marked nuclei, and we examined only initial timepoints since imaging the dyes disrupted development, to assess whether chromosome individuality appeared similar as in fixed, DAPI-stained embryos. First, we used the Hoechst 33342 dye NucBlue. The NucBlue signal was clearly visible only in nuclei closest to the objective lens; the signal became diffuse or undetectable deeper into the embryo, and cytoplasmic foci were bright. Therefore, we also soaked embryos in the newer live-cell DNA dye Kakshine Yellow (Uno *et al*., 2021), which we found worked better for seeing cells deeper into the embryos and produced more specific staining of chromosomes. We found some nuclei with condensed chromosomes akin to what we observed by DAPI staining in fixed samples, which we interpret as likely mitotic stages, but most nuclei stained by either live dye had a wider staining pattern at each chromosome than we saw by DAPI staining (Figure 3A,B); we interpret these as being in interphase. Kakshine dyes and NucBlue fluoresce much more brightly when bound to DNA than when they are bound to RNA (Uno *et al*., 2021), although the extent to which this is true in living *H. exemplaris* embryos is not known. Because we also saw bright foci of signal throughout the cytoplasm of the *H. exemplaris* embryonic cells (Figure 3A,B), we do not know whether the nuclear signal represents only DNA or also RNA and perhaps nascent mRNA associated with chromosomes; in either case, the NucBlue and especially Kakshine Yellow patterns afforded us a chance to observe whether the chromosome individuality we observed by DAPI staining of fixed embryos could also be seen in the absence of fixation. The broad regions of both NucBlue and Kakshine Yellow fluorescence appeared highly individualized (Figure 3A,B), confirming that chromosomes remain unusually individualized in live embryos.

**Figure 3.**
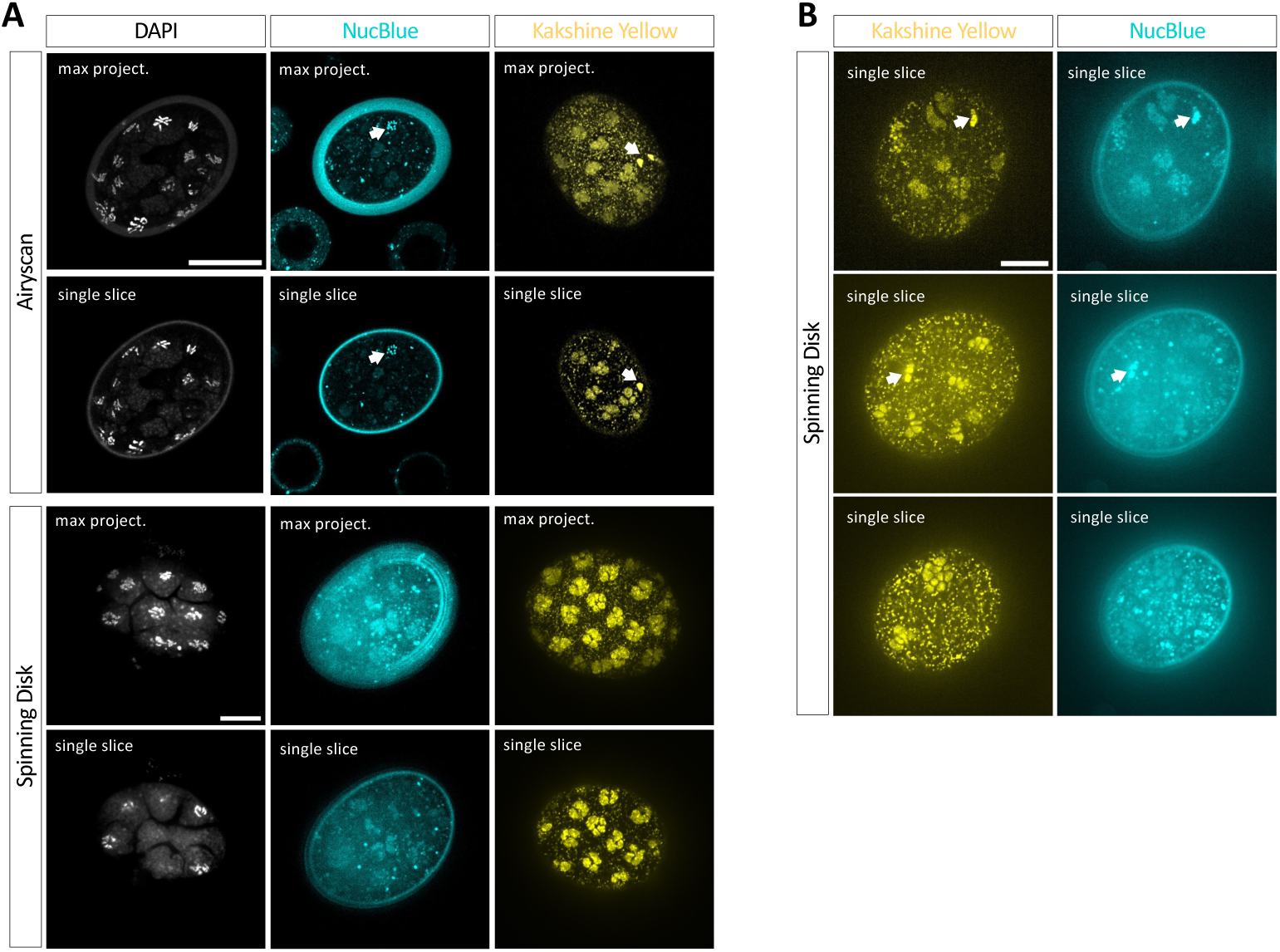
Live DNA staining of *H. exemplaris* early embryos confirms chromosome individualization. (A) Comparison of fixed, DAPI stained embryos with live NucBlue or Kakshine Yellow staining in representative tardigrade embryos, showing single slices or maximum intensity projections (max project.) through the layer of nuclei closest to the objective lens, imaged by Airyscan confocal (top) or spinning disk confocal imaging (bottom). (B) Three representative embryos double-stained with NucBlue and Kakshine Yellow. Arrows point to nuclei that we interpret as mitotic stage nuclei. All scale bars: 20µm.

### Chromosomes appear to be spatially separated from each other in nonspherical *H. exemplaris* early embryonic nuclei

Because none of the staining methods above revealed any signs of typical spherical nuclei, even in background staining patterns, we sought to use other methods to infer the shape of the nucleus. To visualize the overall nuclear shape, we performed *in vivo* staining with Nile Red, which detects neutral lipids including cytoplasmic lipid droplets (Greenspan *et al*., 1985). We expected Nile Red to stain the cytoplasm and outline weakly-fluorescent or nonfluorescent nuclei. As expected, we found that chromosome regions were unstained in all cells of embryos soaked in Nile Red solution at different dye concentrations and soaking times. We also observed weak Nile Red fluorescence between chromosomes (Figures 4A-C), which appeared similar in mitosis (Figure 4A,B) and interphase (Figure 4A,C). We could not detect typical spherical nuclei in the staining patterns of any embryos. We imaged Kakshine Yellow and Nile Red together to see where chromosomes were in relation to the Nile Red pattern. Although there was some expected bleedthrough between fluorescence channels, the Kakshine Yellow chromosome pattern could be seen to mark the Nile Red-free nucleoplasm (Figure 4D).

**Figure 4.**
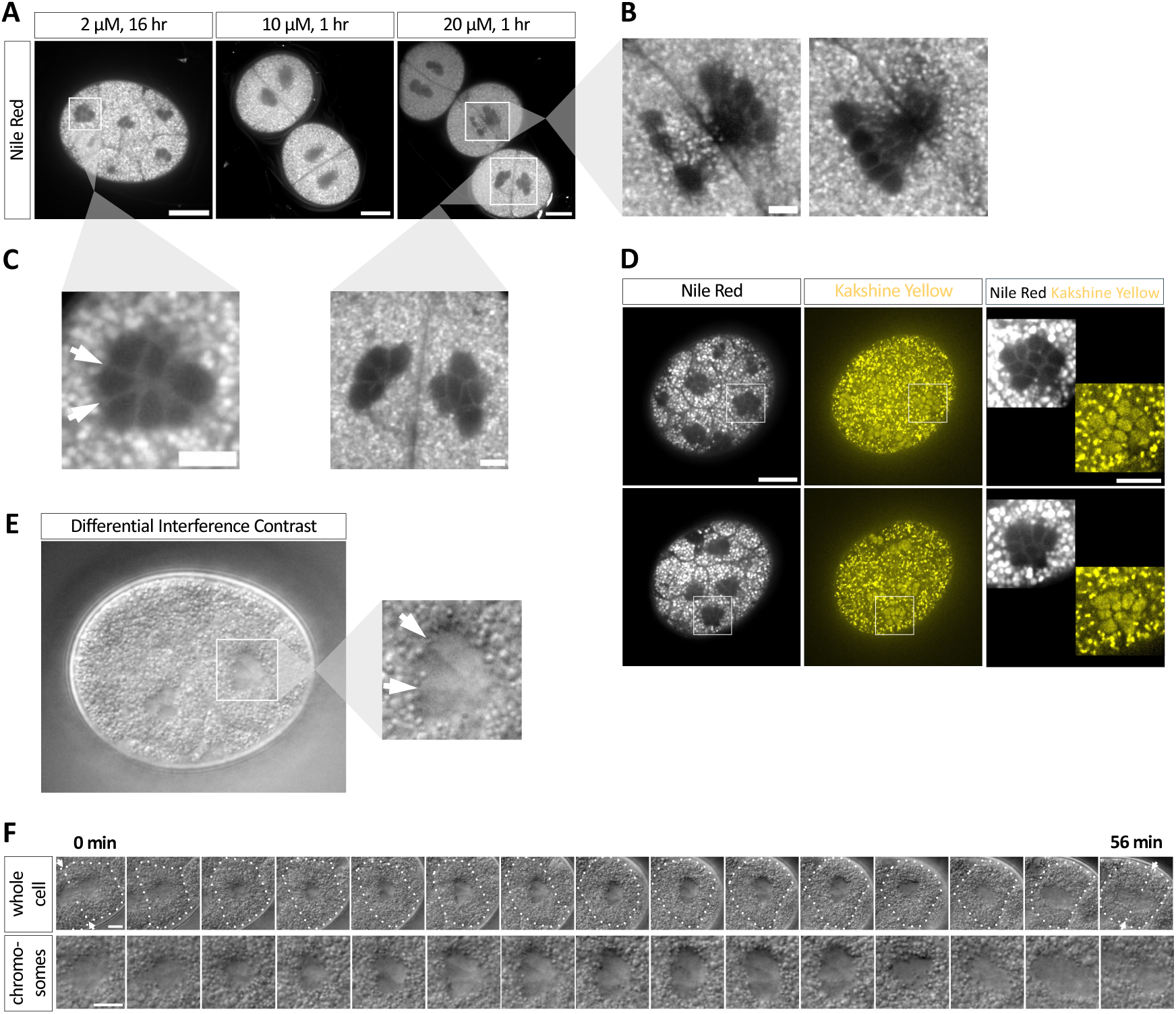
Chromosomes appear to be spatially separated in nonspherical nuclei of *H. exemplaris* early stage embryos. (A) Nile Red-staining enrichment patterns in representative tardigrade embryos at various dye concentrations and staining durations. White box labels the zoomed-in cell shown in Figure 4B. Scale bars, 20 µm. (B) Enrichment patterns of Nile Red-staining in a zoomed-in cell undergoing cell division from Figure 4A, two focal planes. Scale bar, 5 µm. (C) Enrichment patterns of Nile Red-staining in zoomed-in nuclei from Figure 4A. Arrows point to Nile Red-stained divisions between individual chromosomes. Scale bars, 5 µm. (D) Enrichment patterns of Nile Red-staining and Kakshine Yellow-staining in two representative tardigrade embryos, with zoomed-in views of cells in higher contrast at right. Scale bars: 20 µm for whole embryo views, 10 µm for insets at right. (E) DIC image of a 16-cell stage embryo, with a zoomed-in view showing in higher contrast the center of an interphase stage cell. Arrows point to divisions apparent between individual chromosomes. Scale bar, 20 µm. (F) DIC timelapse of the same cell through its entire cell cycle, from the cell division that produced it to its own division (arrows mark division planes), whole cell (top) and zoomed-in view of chromosomes (bottom). Images are shown every 4 minutes for 56 minutes in total. Dotted outline marks the cell of interest. Scale bars, 5 µm.

By differential interference contrast (Nomarski) light microscopy imaging of live embryos, in cells that we could confirm by time-lapse imaging were between mitoses and hence in interphase, we also typically saw non-spherical nuclei with individual chromosomes apparent (Figure 4E,F). These results suggest that chromosomes in *H. exemplaris* are spatially separated from each other, and, since we did not observe a single nucleus encasing all of the chromosomes by any imaging methods, likely within a nuclear form that is atypical of animal cells.

### Nuclear lamina surrounds individual chromosomes in *H. exemplaris* early embryonic cells

Given that chromosomes remain more individualized in *H. exemplaris* early embryos than expected, and that we found a neutral lipid marker separating chromosomes, we next asked whether the nuclear lamina has a typical spherical appearance in these cells, and whether a barrier separates chromosomes from each other. We immunostained embryos for nuclear lamins to examine the morphology of the nuclear lamina. *H. exemplaris* has two nuclear lamin proteins (Hering et al., 2016), and we found that antibodies to one of them, lamin-1, marked regions around chromosomes with minimal cytoplasmic staining. Antibodies to the other lamin, lamin-2, produced bright staining through the cytoplasm in embryos, plus similar staining patterns as anti-lamin-1, but weakly (data not shown). We were surprised to see that the anti-lamin-1 staining surrounded each chromosome (Figure 5, Figure S4). Anti-lamin-1 staining had been shown previously in a roughly spherical shape plus in regions of nucleoplasm in at least some tissues of hatched *H. exemplaris* animals (Hering *et al*., 2016) but lamin-1 localization had not been reported in embryos previously. We observed lamin-1 surrounding each chromosome throughout multiple early embryonic cell cycles, through at least the 32 cell stage (Figure 5C). In nearly every nucleus we examined, looking all of the nuclei visible in the half of each embryo (n = 10) closer to the coverslip, where we could distinguish chromosomes clearly, we saw multiple lamin-bound compartments around chromosomes in nearly every cell (91/92 cells, 98.9%; in one case we saw no anti-lamin staining). We conclude that nuclear lamina surrounds individual chromosomes in these cells. This arrangement is atypical of animal cells, in which the nuclear lamina generally forms a sheet at the periphery of a single, spherical nuclear envelope (Gruenbaum and Foisner, 2015).

**Figure 5.**
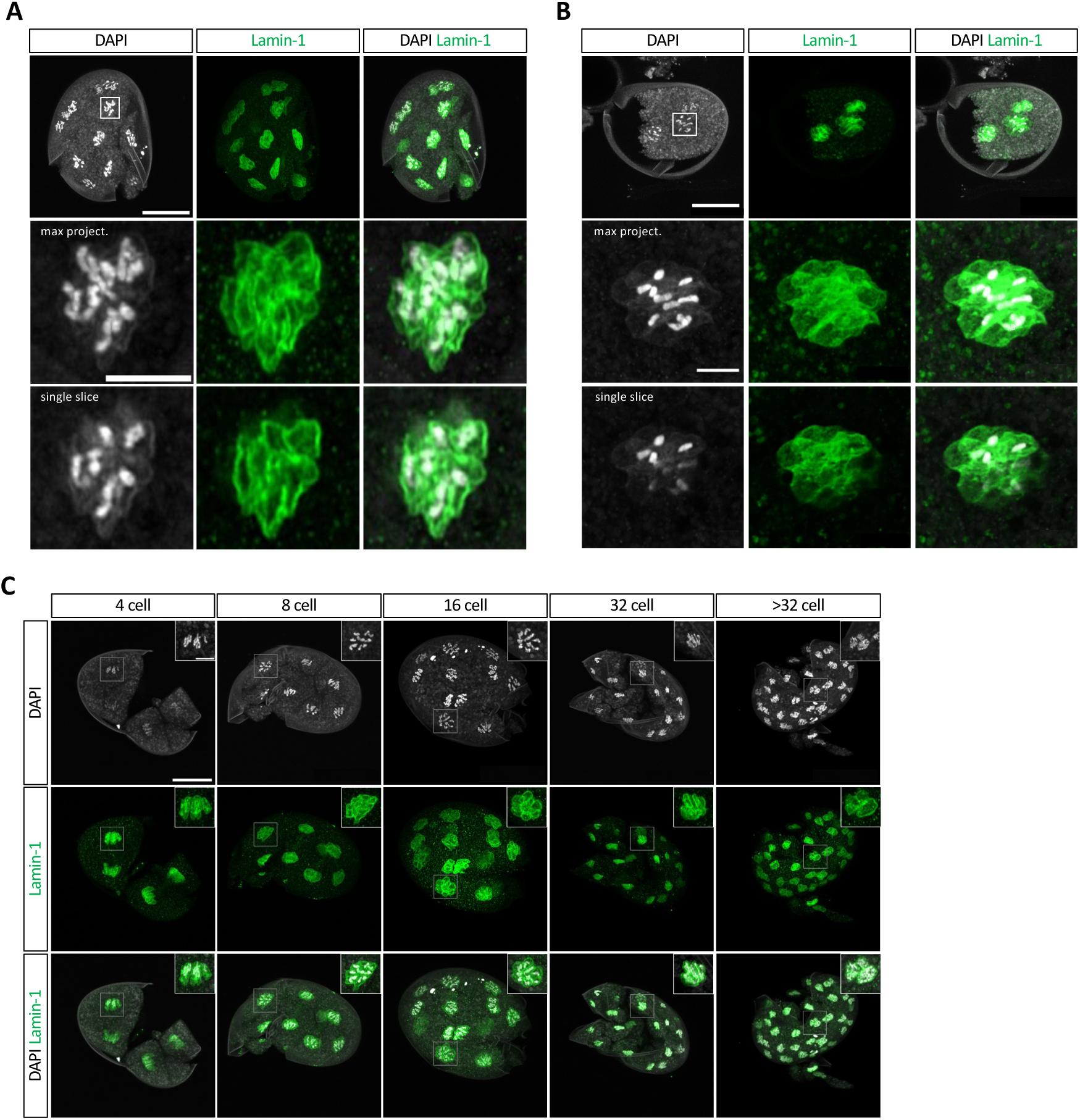
Nuclear lamina form around individual chromosomes creating either fully or partially separate compartments in *H. exemplaris* early embryonic nuclei. (A-B) Localization patterns of DAPI (white) and lamin-1 (green) in representative early *H. exemplaris* embryos. White boxes label the zoomed-in nuclei, showing single slices or maximum projections (max project.) in the images below. Scale bars: 20 µm for whole embryo views, 5 µm for zoomed-in nuclei below. (C) Localization patterns of DAPI (white) and lamin-1 (green) in representative 2 to 32-cell *H. exemplaris* embryos, with zoomed-in views of cells with contrast adjusted in top right corners. Scale bars: 20 µm for whole embryo views, 5 µm for insets. Embryos were imaged using a Zeiss 880 microscope with a fast Airyscan detector, and the individual nuclei in A and B were imaged with a super-resolution Airyscan detector.

## DISCUSSION

We have examined chromosome and nuclear form in early embryos of the tardigrade *H. exemplaris*. We found that DAPI-stained chromosomes in fixed, interphase cells were more condensed than in *C. elegans* embryos, a representative of typical cells, and we were surprised to find that chromosomes remained individualized even through interphase. Staining methods did not reveal any signs of a typically spherical nuclear lamina and envelope surrounding all of the chromosomes in a cell. Instead, we found by anti-lamin staining that each chromosome was either fully or partially surrounded by lamin. These results reveal that chromosome and nuclear architecture in the early embryos of this tardigrade species are highly unusual, with chromosomes packaged separately instead of entangled with each other in interphase.

We found that chromosomes appeared more condensed in fixed, DAPI-stained embryos than in live, NucBlue-or Kakshine Yellow-stained embryos. However, both NucBlue and Kakshine Yellow also stained cytoplasmic foci brightly, much more so than would be expected if the cytoplasmic signals were mitochondrial DNA alone. We speculate that NucBlue and Kakshine Yellow might also stain RNA in living *H. exemplaris* embryos more brightly than was measured previously in other contexts (Uno et al., 2021), and/or that the concentration of RNA is especially high in places including nascent mRNAs such as around chromosomes. Consistent with this idea, RNA-Seq of *H. exemplaris* embryos has produced mRNA abundance patterns that suggest that the early embryos are actively transcribing genes (Levin et al., 2016). Given the likelihood that NucBlue and Kakshine Yellow brightly marked more than just the DNA, we used DAPI staining in parallel in *H. exemplaris* and *C. elegans*, under the same conditions, to make quantitative comparisons of condensation states. We also confirmed the *H. exemplaris* staining pattern we saw using an antibody recognizing acetylated histone H4. Taken together, these results suggested that interphase chromosomes in early tardigrade embryos are more condensed than in *C. elegans*, yet are unlikely to be composed primarily of transcriptionally silenced heterochromatin. This condensation may have relevance to extremotolerance biology. Past studies from other organisms found that anoxia can transiently induce signs of chromosome condensation in certain organisms including *Drosophila*, *C. elegans*, and zebrafish (Foe and Alberts, 1985; Padilla and Roth, 2001; Padilla et al., 2002), and condensed chromosomes have more resistance to ionizing radiation than decondensed chromatin (Suzuki et al., 2009; Takata et al., 2013). Condensation might also keep ends resulting from double-strand DNA breaks close together, allowing NHEJ to accurately ligate broken ends in the face of many simultaneous double-strand breaks. Resolving the precise degree of chromatin condensation in living tardigrade embryos will be of interest for understanding the degree to which the chromatin has atypical organization.

All of the methods above, including live staining of nucleic acids, also revealed that chromosomes were highly individualized in *H. exemplaris* embryos. Chromosomes have been found to be transiently individualized in karyomeres, i.e. in separate nuclear envelopes, in telophase of some animals with large cells, such as the embryos of certain amphibians, polychaete worms, sea snails, or sea urchins (Conklin, 1902; Longo, 1972; Emanuelsson, 1973; Montag et al., 1988; Lemaitre et al., 1998). The structures we observed resemble such karyomeres, except that karyomeres are temporary structures; they fuse with each other soon after forming, during telophase, and interphase nuclei appear normal in these organisms. We speculate that mechanisms that fuse karyomeres in other organisms may be absent or inhibited in *H. exemplaris* early embryos.

At later embryonic stages, we did not observe *H. exemplaris* chromosomes in a more condensed state than in *C. elegans*, nor in at least some adult tissues, although typical resolution limits of light microscopy might have made condensation state difficult to detect in small cells. It is possible that electron microscopy will be required to resolve similar features and perhaps other unexpected features of chromosome packaging in such small cells, and electron microscopy might also help further resolve the differences we saw using DAPI in fixed cells vs. Kakshine Yellow or NucBlue in living cells. Electron microscopy would also be valuable for resolving whether, in early embryos, each chromosome is fully enclosed in its own micronucleus, or alternatively whether all chromosomes are surrounded by an unusual nucleus with multiple, connected lobes. We have attempted to collaboratively process *H. exemplaris* embryos for transmission electron microscopy using high pressure freezing, as a step toward focused ion beam scanning electron microscopy for examining 3D structure at high resolution, but we have not yet found satisfactory processing methods. Hering et al (2016) observed more typical anti-lamin patterns in at least some cells of hatched *H. exemplaris* animals, and oogenic nuclei appeared typical in electron microscopy images of oocytes (Jezierska et al. 2021), so the unusual features we observed are not present in all *H. exemplaris* cells.

It is not known if the unusual features we observed in embryos exist in tardigrades more generally, or alternatively in just this species – and in the small number of other animal groups that can survive desiccation with loss of nearly all intracellular water, for example rotifers (see Crowe et al., 1992 for review). We have attempted similar imaging in some other tardigrade species, but the ornamented eggshells of other tardigrade species (see Bertolani et al., 1996 for review) presented challenges for imaging embryos, and we have not yet succeeded in attempts to minimize this problem by matching refractive index of media to the eggshells’ refractive index. We had initially chosen *H. exemplaris* as a model in part because their embryos have smooth eggshells, so that imaging through the eggshell would not pose problems (Gabriel et al., 2007; Goldstein, 2022).

In typical animal cells, chromosomes are highly entangled in interphase and are decatenated (untangled) by topoisomerase enzyme activity (Coelho et al., 2003), and also dependent on an RNA-binding domain-containing protein, SRBD1 (Raaijmakers et al., 2025). Mitotic chromosomes maintain individuality via proteins that can present a steric and electrostatic charge barrier to chromosome contacts (Cuylen et al., 2016). Our finding that lamin-1 fully or partially surrounds each chromosome in early *H. exemplaris* embryos suggests that these cells use some distinct mechanisms from those above to maintain chromosome individuality. We have not yet found a method that will label the nuclear membrane in embryos (see Materials and Methods), which would be valuable for determining whether chromosomes surrounded by lamin are enclosed in separate nuclear envelopes. *H. exemplaris* lamin-1 lacks a CAAX domain and so might not associate directly with nuclear membrane, but lamin-2 has a CAAX domain (Hering et al., 2016). Recent efforts to develop CRISPR and transgenic methods for species of tardigrades (Kumagai et al., 2022; Tanaka et al., 2023; Kondo et al., 2024) may make it possible to fluorescently tag these lamins and nuclear envelope proteins to help further resolve questions about nuclear structure, as well as to resolve whether even modest condensation and decondensation cycles occur in the embryonic stages we observed.

Our finding that chromosomes remain individualized throughout the cell cycle in *H. exemplaris* embryos may have evolutionary implications, given what is already known about how these animals survive DNA damaging conditions. We found previously that *H. exemplaris* subjected to DNA-damaging gamma radiation, or to a radiomimetic drug, upregulated base excision repair and NHEJ genes to a remarkable degree, and other labs have seen similar results in this species and in a second species in the same genus (Clark-Hachtel et al., 2024; Anoud et al., 2024; Li et al., 2024). Base excision repair mechanisms make precise repairs to damaged bases on otherwise unbroken double stranded DNA. NHEJ repairs DNA double-strand breaks, and is error prone, joining broken ends and even blunt ends, without relying on a homologous template for accurate repair. Desiccation induces DNA double strand breaks, as does radiation (Mattimore and Battista, 1996), and we predict that tardigrades must have mechanisms that maintain chromosome number in the face of many, simultaneous DNA breaks during desiccation events in nature. Repair of multiple DNA double strand breaks simultaneously after desiccation by NHEJ would be expected to result in many incorrect ligations of broken DNA ends. Yet Hi-C assemblies of two different *Hypsibius* species showed significant synteny and only minimal between-chromosome rearrangements (just one chromosome fusion or break was found, and no apparent translocations; Li et al., 2024; Hoencamp et al., 2021). We speculate that the atypical chromosome organization we observed, with chromosomes kept separate from each other through interphase in early embryos, might contribute to limiting translocations and chromosome fusions in the face of many DNA breaks during natural desiccation events.

## MATERIALS AND METHODS

### Maintenance of Cultures/Strains

Cultures of *Hypsibius exemplaris* (Z151) were maintained as previously described (McNuff, 2018). Tardigrades were kept in 35 mm vented petri dishes (Tritech Research, T3500) with approximately 2 mL of spring water (Deer Park) and 0.5 mL Chloroccocum sp. algae (Carolina Biological Supply). Cultures were split approximately once a month and kept in a plastic box with wet paper towels lining the bottom to maintain humidity.

*Caenorhabditis elegans* (N2 strain) was maintained on Nematode Growth Medium plates at 20°C under standard conditions (Brenner, 1974).

### Fixed DNA Staining of Embryos

Fixed DNA staining to visualize chromosomes in early tardigrade embryos was performed following the fixation steps in published methods (Smith and Gabriel, 2018). Gravid adults were collected and the time was recorded when a clutch of embryos was laid. The embryos were then cut out of the cuticle and fixed with 4% paraformaldehyde at specific time intervals. After fixation, the eggshells were nicked and the embryos were mounted onto slides with 2 mL DAPI fluoromount-G (Southern Biotech) with 28.41 µm glass microspheres (Whitehouse Scientific’s Monodisperse Standards). Nicking often results in damage to parts of embryos; only embryos with minimal damage, away from areas of interest, were examined. The protocol was repeated until there were representative embryos for every 15 minutes from 2 to 6hpl. A total of 61 embryos were imaged with an average of about 3 embryos per time point. The timeline covered the 2 to 32 cell stages in tardigrade embryos because a typical cell cycle in *H. exemplaris* is about 50 minutes after the first division, which lasts 2 hours (Gabriel *et al*., 2007). Cell divisions are not synchronous (Gabriel *et al*., 2007), but for convenience, we grouped and named embryos in figures by the stage that had been reached at the previous complete round of division for all cells (for example, we use “8 cells” to refer to all embryos that reached the 8-cell stage but had not yet reached the 16-cell stage).

The fixed DNA staining protocol was repeated in *C. elegans* embryos for a species comparison. Embryos were dissected from gravid adults ranging from the 2 to 16 cell stage. Both *H. exemplaris* and *C. elegans* embryos were fixed in the same tube and imaged on the same slides to ensure identical conditions. All embryos were imaged with a 60x oil objective at the same settings using a Zeiss 880 with fast Airyscan detector.

### Fixed DNA Staining of Adults

Gravid adults were collected after they laid embryos and crawled out of the cuticle to reduce autofluorescence of the gut. Animals were anesthetized with 20 mM levamisole (L9756, MilliporeSigma) for 30 minutes before fixation. Fixation was performed with 4% paraformaldehyde in phosphate-buffered saline and 0.1% TritonX for 30 minutes at room temperature. The fixative was washed out with 0.5x PBT. After fixation, the adults were mounted onto slides with 2 mL DAPI fluoromount-G (Southern Biotech) with 28.41 µm glass microspheres (Whitehouse Scientific).

The fixed DNA staining protocol was repeated in *C. elegans* adults for a species comparison. All adults were imaged with a 60x oil objective at the same settings using a Zeiss 880 with fast Airyscan detector.

### Live Imaging of Embryos

Live DNA staining to visualize chromosomes in early tardigrade embryos was attempted with 33 different DNA dyes (in alphabetical order): Acridine Orange, DRAQ5 Fluorescent Probe Solution (ThermoFisher), Kakshine Orange (Cosmo Bio Co.), Kakshine Red (Cosmo Bio Co.), Kakshine Yellow (Cosmo Bio Co.), NucBlue Live ReadyProbes Reagent (Hoechst 33342, ThermoFisher), NucleoLIVE (Saguaro Biosciences), NucRed Live ReadyProbes Reagent (ThermoFisher), Propidium Iodide, SiR-DNA (Spirochrome), SPY505-DNA (Spirochrome), SPY650-DNA (Spirochrome), SYTO Green Fluorescent Nucleic Acid Stains (SYTO 11 through 14, 16, 21, 24, 25, ThermoFisher), SYTO Orange Fluorescent Nucleic Acid Stains (SYTO 80 through 85, ThermoFisher), and SYTO Red Fluorescent Nucleic Acid Stains (SYTO 17, 59 through 64, ThermoFisher). The two dyes that penetrated eggshells (i.e., showed some staining in embryos) and showed the most specific staining of chromosomes were NucBlue and Kakshine Yellow. We also tried endoplasmic reticulum and golgi staining by ER Tracker Red (ThermoFisher BODIPY TR Glibenclamide) and Cytopainter ER/Golgi Staining Kit (AbCam 139485), but neither worked for us on intact embryos.

For NucBlue staining, two to three drops were added to 500 mL of spring water in a 1.5 mL tube, and embryos were cut out of the cuticle they were laid in and placed in the reagent in a 35 mm vented petri dish or a glass depression slide to soak for a range of 30 minutes to 2 hours. The dish or slide was kept in the dark in a tinfoil covered humid box at room temperature. After soaking, the embryos were mounted onto slides with spring water with 28.41 µm glass microspheres (Whitehouse Scientific) and the coverslip was sealed with melted Valap (1:1:1 mixture of Vaseline, lanolin, and paraffin) maintained at 70°C and applied with a paintbrush. All embryos were then immediately imaged with the same settings using a Zeiss 880 with fast Airyscan detector or a Yokogawa CSU-X1 spinning disk confocal (Nikon, BioVision).

For Kakshine Yellow (PC1, Uno et al., 2021) staining, 1µL staining solution was added to 500µL of spring water in a 1.5mL tube, and then the entire solution was added to a glass depression slide, which was then placed into a foil-covered humid chamber. Embryos from the same clutch were cut out of the cuticle they were laid in and placed in the reagent to soak for 2hrs, rocking gently at room temperature. After soaking, the embryos were mounted onto slides with spring water with 28.41 µm glass microspheres (Whitehouse Scientific) and the coverslip was sealed with Valap. All embryos were then immediately imaged with the same settings using a Zeiss 880 with fast Airyscan detector or a Yokogawa CSU-X1 spinning disk confocal (Nikon, BioVision), both with a 60x oil objective. Maximum intensity projections encompassed the entirety of the layer of nuclei visible closest to the objective (usually between 6 and 15µm in height).

For Nile Red staining, solutions of Nile Red (Invitrogen N1142) from 2µm to 20µm were prepared in spring water, and embryos and adults were soaked together for 1-16 hours, followed by three 5-minute washes and a 15-minute wash in 20 µM levamisole in spring water. Embryos were then mounted onto a slide in spring water together with 12-18 µm glass microspheres (Whitehouse Scientific), and the coverslip was sealed with Valap. Embryos were then imaged on a Yokogawa CSU-X1 spinning disk confocal (Nikon, BioVision) using a 60x oil objective.

DIC imaging used a Nikon E800 upright microscope with a 60x oil objective and a Photometrics CoolSnap Dyno camera.

### Immunostaining of Embryos

Immunostaining in early tardigrade embryos was performed following published methods (Smith and Gabriel, 2018). Lamin immunostaining to visualize the nuclear envelope was conducted using the anti-lamin-1 primary antibody generated by the Mayer Lab (Hering *et al*., 2016). Embryos ranging from 2 to 6 hpl were collected. The immunostaining was performed using the anti-lamin-1 antibody at a concentration of 1:250. Goat anti-guinea pig::Alexa Fluor 488 (A11073, ThermoFisher) secondary antibody at a concentration of 1:500 was used to detect the primary antibody. Immunostaining to detect histone H4 acetylation was performed using an anti-histone H4ac (pan-acetyl) primary antibody (AB_2793201, Active Motif) at a concentration of 1:500. Donkey anti-rabbit::Alexa Fluor 488 (A32790, Invitrogen) secondary antibody at a concentration of 1:500 was utilized to detect the primary antibody. Embryos were mounted onto slides with 2 mL DAPI fluoromount-G (Southern Biotech) with 28.41 µm glass microspheres (Whitehouse Scientific). The embryos were imaged using a Zeiss 880 microscope with the fast Airyscan detector, and individual nuclei were imaged with the super-resolution Airyscan detector.

### Image Quantification

Chromosome condensation in both *H. exemplaris* and *C. elegans* embryos was quantified using a previously developed fluorescence-based condensation assay (Maddox *et al*., 2006). For each species, 2-5 nuclei per embryo for 20 embryos were analyzed. The number of nuclei varied due to the different cell stages of the embryos. The first 2-5 nuclei that were closest to the cover slip when going through the z-stack, specifically the first 2-5 interphase nuclei for *C. elegans* embryos, were chosen for analysis to create consistent criteria. Maximum intensity projections were generated for each nucleus using ImageJ. The smallest square/rectangular region that fit the nucleus, or encapsulated all the chromosomes for *H. exemplaris*, was exported as a text file, where each pixel was represented as a number. The remainder of the quantification was done in Excel. The text files of the nuclei were individually scaled to set the minimum intensity to 0 and the maximum to 255 so the fluorescence intensity was independent of variation from photobleaching. Histograms of the fluorescence intensity from 0 to 255 for each nucleus were then created. From that data, graphs of the percentage of pixels below thresholds set at 20%, 35%, and 50% of the maximum intensity (values of 51, 89, and 127, respectively) were also constructed in SuperPlotsofData (https://huygens.science.uva.nl/SuperPlotsOfData/). The mean and 95% confidence intervals were conservatively determined, treating each embryo as a replicate (n=20). Statistical comparisons used Welch’s t-test.

### Imaris Segmentation Machine Learning

Chromosome segmentation in *H. exemplaris* embryos was done using Imaris image analysis workstation. Machine Learning using the Surface model was trained on 5 raw DAPI images for each cell stage from 2 to 32-cell embryos. The learning software was then run on a batch of raw DAPI images that were captured for the comprehensive timeline of fixed, DNA-stained embryos. Once the segmentation had run, the resultant images were individually analyzed to circle each nucleus and label it as a class. Statistical data that were automatically calculated within Imaris for each class were exported to Excel. The number of labelled surfaces (chromosomes) per class (nucleus) was used to create the histogram in Figure 2B representing the number of chromosomes segmented in each nucleus within the batch of embryos.

## ACKNOWLEDGMENTS

We thank the members of the Goldstein Lab for support and feedback on this project, including Kira Heikes for training and Courtney Clark-Hachtel for sharing previously unpublished results that motivated this research. We thank Lars Hering and Georg Mayer for sharing anti-lamin antibodies, Yoshikatsu Sato for sharing Kakshine dyes, Nat Prunet for assistance in the UNC Biology microscopy core, Paul Maddox for advice implementing his quantitative methods for assessing chromosome condensation, and Jenny Tenlen, Lars Hering, and Georg Mayer for comments on the manuscript. This work was supported by a grant from the National Science Foundation (IOS 2028860 to B.G.), a UNC Chapel Hill NIH Training Grant in Genetics (T32GM135128, supporting A.N.C.), and a grant from the Office of Undergraduate Research at UNC via the Summer Undergraduate Research Fellowship (to L.P.).

---

**Figure S1.**
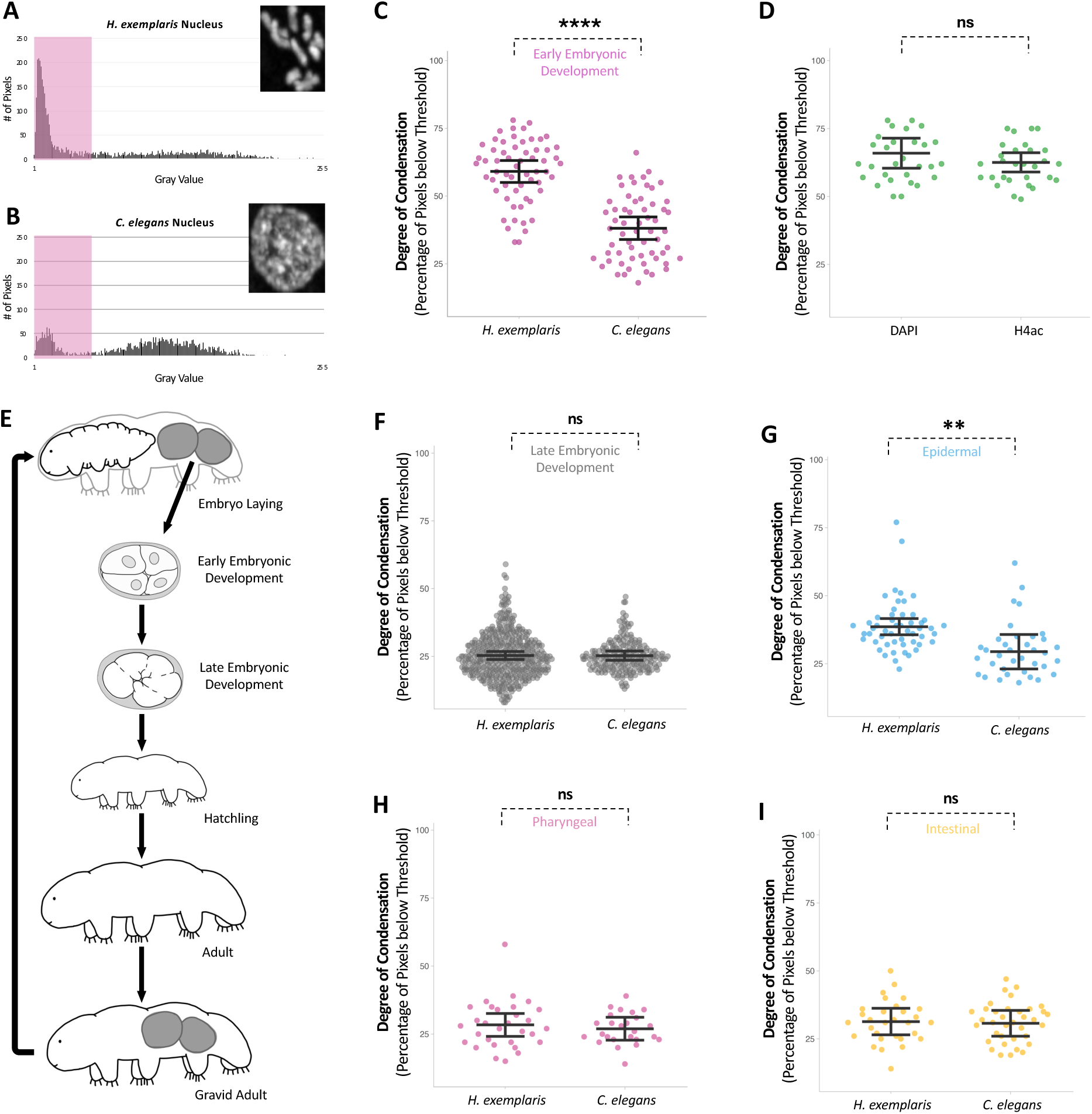
Chromosome condensation quantification throughout *H. exemplaris* and *C. elegans* early embryonic development, late embryonic development, and adult life stages. (A) Representative graph of the fluorescence intensity distribution in one *H. exemplaris* nucleus. The section highlighted is the region below 20% of the maximum intensity threshold (below 51 gray value). (B) Representative graph of the fluorescence intensity distribution in one *C. elegans* nucleus. The section highlighted is the region below 20% of the maximum intensity threshold (below 51 gray value). (C) Plot comparing the percentage of pixels below 20% of the maximum intensity threshold for all *H. exemplaris* (n = 20) and *C. elegans* (n = 20) early developmental embryos. Bars represent the mean and 95% confidence intervals. \*\*\*\**P* < 0.0001. (D) Plot comparing the percentage of pixels below 20% of the maximum intensity threshold for all *H. exemplaris* embryos with DAPI (n = 7) and histone H4ac staining (n = 7). Bars represent the mean and 95% confidence intervals. (E) Schematic diagram of the tardigrade *H. exemplaris* life cycle. (F) Plot comparing the percentage of pixels below 20% of the maximum intensity threshold for all *H. exemplaris* (n = 43) and *C. elegans* (n = 20) late developmental embryos. For *H. exemplaris*, this covered the 60 cell stage to hatchling. (G) Plot comparing the percentage of pixels below 20% of the maximum intensity threshold for epidermal nuclei in *H. exemplaris* (n = 10) and *C. elegans* (n = 7) adults. (H) Plot comparing the percentage of pixels below 20% of the maximum intensity threshold for pharyngeal nuclei in *H. exemplaris* (n = 10) and *C. elegans* (n = 5) adults. (I) Plot comparing the percentage of pixels below 20% of the maximum intensity threshold for intestinal nuclei in *H. exemplaris* (n = 10) and *C. elegans* (n = 7) adults. Related to Figure 1.

**Figure S2.**
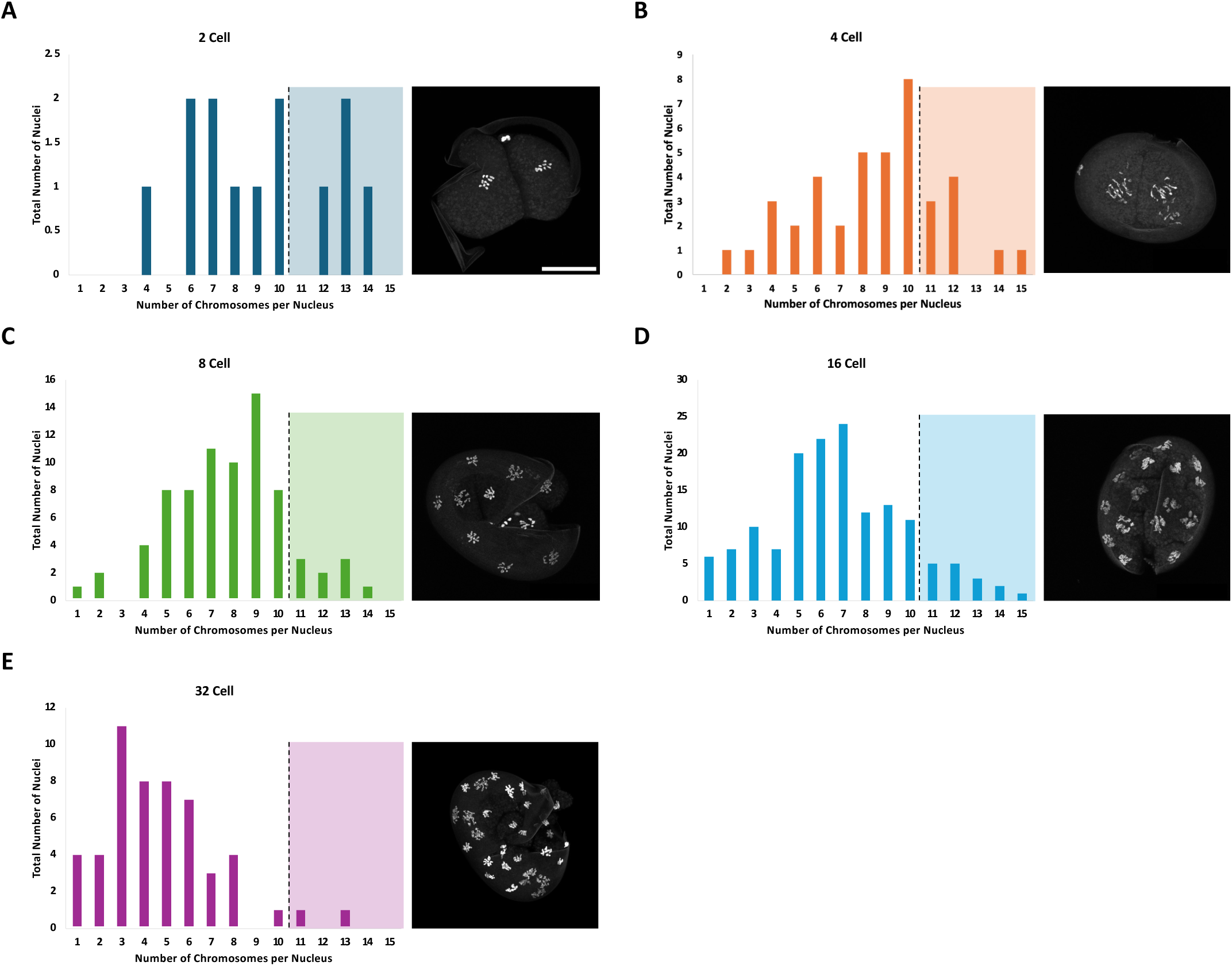
Histograms of the chromosome counts for the (A) 2 cell, (B) 4 cell, (C) 8 cell, (D) 16 cell, and (E) 32 cell stages in *H. exemplaris* early embryos. Representative images of embryos at each cell stage are to the right of the corresponding graphs. Scale bar, 20 µm. The section colored on the right side in each graph is the chromosome counts greater than the known number in *H. exemplaris*, 2n = 10 chromosomes. Related to Figure 2.

**Figure S3.**
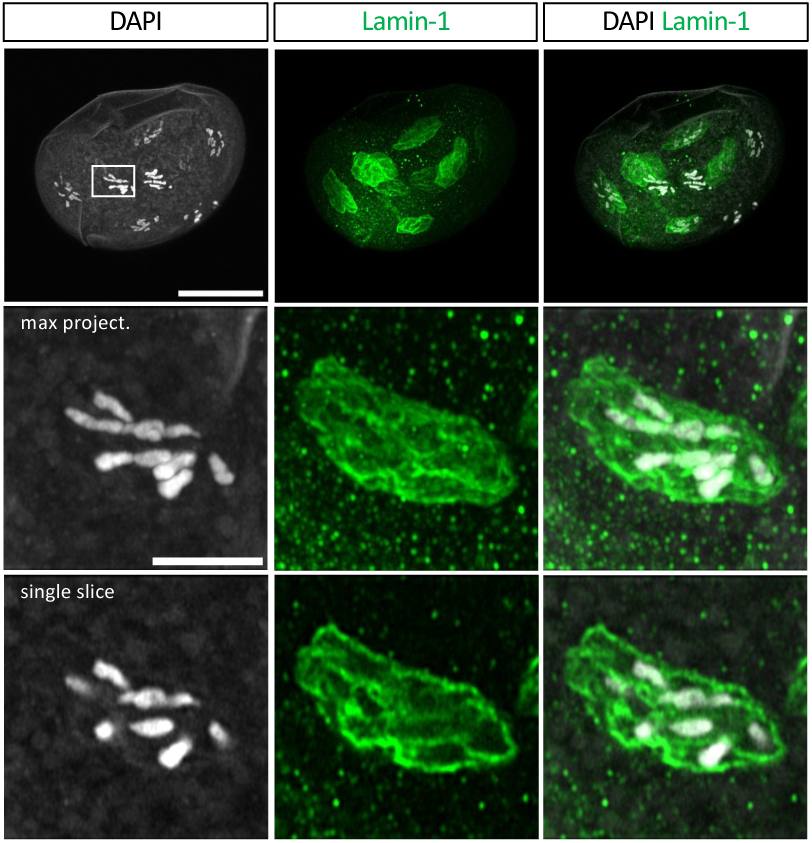
Further example of nuclear lamin-1 staining in *H. exemplaris* early embryonic embryos. White boxes label the zoomed-in nuclei, showing single slices or maximum projections (max project.) in the images below. Scale bars: 20 µm for whole embryo views, 5 µm for zoomed-in nuclei below. Embryos were imaged using the Zeiss 880 microscope with the fast Airyscan detector, and the individual nucleus was imaged with a super-resolution Airyscan detector. Related to Figure 5.

**Supplemental Video**. Videos moving through the full z-stack of representative early *H. exemplaris* embryos from Figure 5C (8 cell, 16 cell, 32 cell, >32 cell) to visualize localization patterns of DAPI (white) and lamin-1 (green). Scale bar = 20 µm. Related to Figure 5.

